# Distinct roles of cohesin-SA1 and cohesin-SA2 in 3D chromosome organization

**DOI:** 10.1101/166264

**Authors:** Aleksandar Kojic, Ana Cuadrado, Magali De Koninck, Miriam Rodríguez-Corsino, Gonzalo Gómez-López, François Le Dily, Marc A. Marti-Renom, Ana Losada

## Abstract

In addition to mediating sister chromatid cohesion, cohesin plays a central role in DNA looping and segmentation of the genome into contact domains (TADs). Two variant cohesin complexes that contain either STAG/SA1 or SA2 are present in all cell types. Here we addressed their specific contribution to genome architecture in non-transformed human cells. We found that cohesin-SA1 drives stacking of cohesin rings at CTCF-bound sites and thereby contributes to the stabilization and preservation of TAD boundaries. In contrast, a more dynamic cohesin-SA2 promotes cell type-specific contacts between enhancers and promoters within TADs independently of CTCF. SA2 loss, a condition frequently observed in cancer cells, results in increased intra-TAD interactions, likely altering the expression of key cell identity genes.

---

Several human cancers, including bladder cancer, Ewing sarcoma and acute myeloid leukaemia, have been associated to prevalent mutations in the *STAG2* gene encoding cohesin subunit SA2 (reviewed in (*1*)). Current evidence suggests that the contribution of cohesin dysfunction to tumourigenesis is not related to cohesion defects or genome instability (*2*-*4*). Instead, it could be the result of altered gene regulation (*5*, *6*). Somatic vertebrate cells carry a second cohesin complex in which SMC1, SMC3 and RAD21 associate with SA1 instead of SA2 (*7*). Cohesin-SA1 most likely performs the essential functions of the complex in *STAG2* mutated cancer cells (*8*). However, it remains unclear to which extent the two variant complexes are functionally equivalent. We therefore set out to characterize the specific roles of cohesin-SA1 and cohesin-SA2 in chromatin architecture in human cells. Since unbalanced amounts of the two complexes could hinder our functional analyses, their relative abundance was assessed in nine human primary cell lines that represent different tissues and lineages (Fig. S1A). Human mammary epithelial cells (HMECs) had high cohesin expression with comparable levels of chromatin-bound cohesin-SA1 and cohesin-SA2. We, thus selected these cells to analyse the genomic distribution of SMC1, SA1 and SA2 by chromatin immunoprecipitation followed by deep sequencing (ChIP-seq). Custom-made, validated antibodies and high depth sequencing (about 100 million reads) ensured whole genome coverage (Table S1). We found that a large fraction of cohesin positions (42,475) were occupied by either variant complex and colocalized with CTCF (Fig. 1A). These “common” positions were featured by high cohesin occupancy and similar read density for SA1 and SA2 (Fig. S1B). In contrast, 39,061 sites in the genome with no CTCF were occupied by SMC1 and SA2 but none or very little SA1 (SA2-only positions hereafter). There was also a small number of cohesin positions (3,198) in which SA1 was the preferred variant with most of them also occupied by small traces of CTCF (SA1-only positions in Fig. 1A, lower right). Analysis of the distribution of these cohesin binding sites in chromatin states defined by ChromHMM (*9*) in HMECs revealed that most SA2-only cohesin positions (77%) were in enhancers, particularly active ones (Fig. 1B). The distribution of the common positions was very different, with only 35% present in enhancers while another 41% were in insulators characterized by the sole presence of CTCF. Half of the SA1-only positions were in heterochromatin, 23% in insulators and only 10% in enhancers. Motif discovery analysis using MEME showed that both common and SA1-only positions were significantly enriched in CTCF binding motif while SA2-only positions were populated by recognition motifs of several transcription factors other than CTCF (Fig. 1C). We validated our findings in MCF10A cells, a non-tumourigenic epithelial breast cell line that, unlike HMECs, can be easily grown and transfected for functional analyses (Fig. 1D and S1C). Also in this case, cohesin SA2-only positions were enriched in enhancers and depleted in insulators (Fig. 1E). The existence of a cohesin-SA2-only population present at non-CTCF sites was further confirmed in a cell line of different embryonic origin (HCAEC, endothelial) in which the number of SA1-only and SA2-only positions was comparable (Fig. 1F and S1C). As in the other cell lines, common and most SA1-only positions overlapped with CTCF. Importantly, a large fraction of common positions was conserved between the epithelial and endothelial cells, whereas SA2-only positions were not, suggesting a role for this cohesin variant in tissue specific transcription (Fig. S1D). Consistent with this possibility, cohesin-SA2 only sites were particularly enriched in superenhancers defined in HMECs (Fig. 1G), which control cell identity genes (*10*). Altogether, our results indicate that cohesin-SA1 function is almost exclusively linked to CTCF, whereas a significant fraction of cohesin-SA2 binds to cis regulatory elements that likely contribute to establish cell type specific transcription programs independently of CTCF.

**Figure 1.**
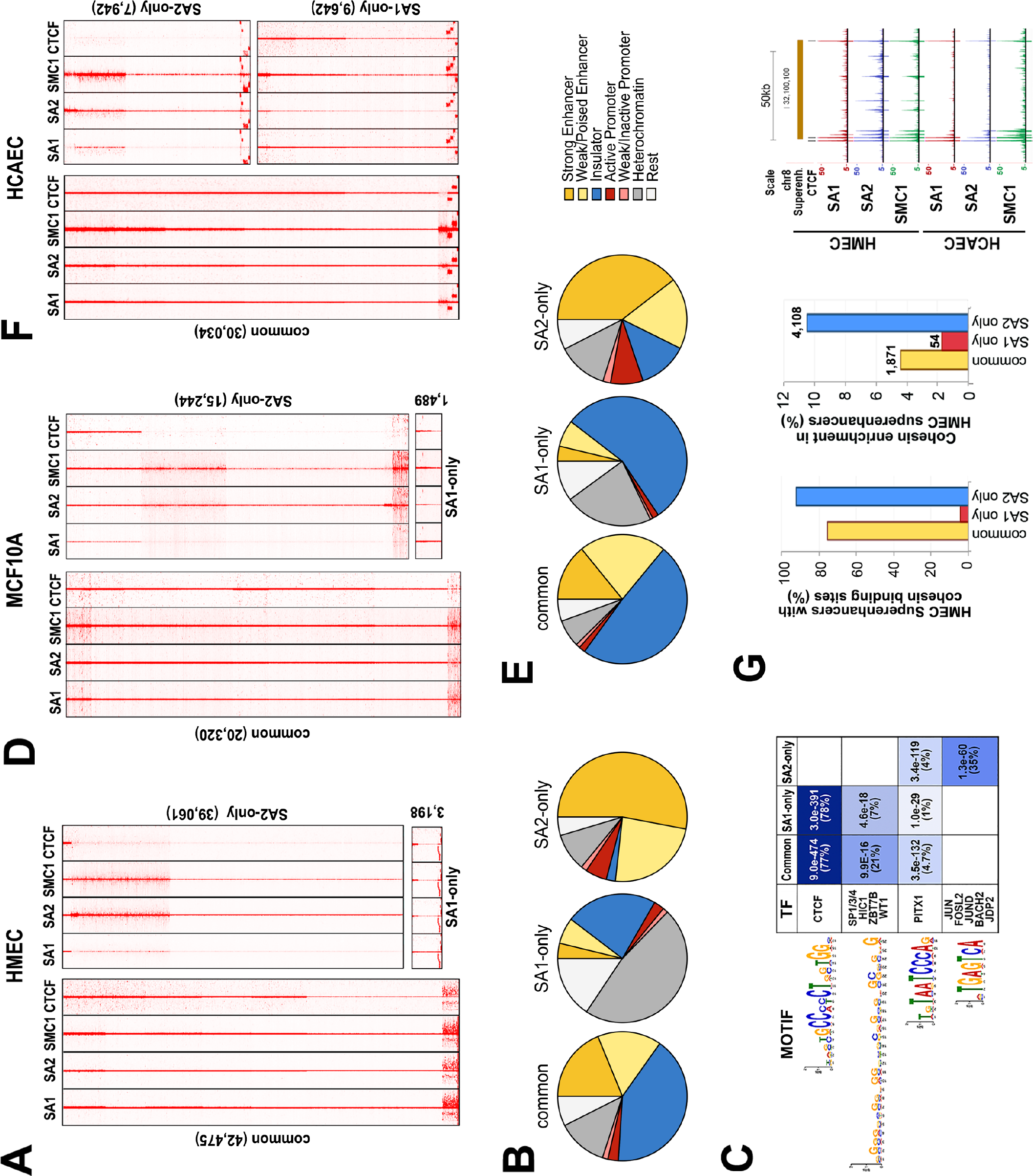
A large fraction of cohesin-SA2 localizes to enhancers independently of CTCF. (A) Analysis of ChIP-seq read distribution for SA1, SA2, SMC1 and CTCF around common, cohesin SA1-only and cohesin SA2-only positions within a 10-kb window in HMECs. (B) Pie charts showing the distribution of cohesin positions in chromatin states defined in HMECs. (C) Motif-based analysis of the indicated positions. E-values of the significantly enriched motifs and percentage of regions containing each motif for a given condition are indicated. “TF” column contains the transcription factors with statistically significant binding to the identified motifs. (D) and (E) Same analysis as in (A) and (B), respectively, for MCF10A cells. (F) Cohesin distribution in human cardiac endothelial cells (HCAECs). (F) Cohesin enrichment in superenhancers defined in HMECs. In the three cell lines, All CTCF datasets are from ENCODE (Table S1).

To further understand how cohesin-SA2 may perform its function, we analysed immunoprecipitates obtained with SA1 and SA2 antibodies from MCF10A cell extracts by mass-spectrometry. Several transcription factors associated with SA2 and not SA1, including Zmym2 (Table S2A). This protein has been described as part of the CoREST transcriptional repressor complex (*11*) and four of its components (HDAC1, HDAC2, KDM1A and RCOR1/3) were also present in the SA2 immunoprecipitates obtained from HeLa nuclear extracts prepared in a way that preserves better transient protein-protein interactions (Table S2B). Coimmunoprecipitation experiments validated the association between SA2 and CoREST and further revealed that most of Zmym2 protein present in the extract was bound to SA2 (Fig. 2A). ChIP-seq analyses confirmed the presence of SA2, and not SA1 or CTCF, at most Zmym2 positions (Fig. 2B). It is therefore possible that the specific interaction of cohesin-SA2 with repressive complexes such as CoREST contributes to gene regulation. To assess the impact of cohesin-SA1 and cohesin-SA2 on gene expression, we transfected MCF10A cells with siRNAs against cohesin subunits and CTCF. Comparable extent of depletion of SA1 or SA2 left similar amounts of cohesin (SMC1) in the cells (Fig. 2C). Using a stringent criterion on RNA sequencing (RNA-seq) data analysis, 157 and 698 differentially expressed genes (DEGs) were identified in cells treated with SA1 and SA2 siRNAs, respectively (Fig. 2D and Table S3-S5). Out of 612 genes deregulated only after SA2 depletion, 417 were not affected by CTCF knock down confirming a CTCF independent role for SA2 in the control of gene expression. Within this group, we found BDNF (brain derived neurotrophic factor), a known target of CoREST in non-neuronal cells (*12*)(Table S3). Gene set enrichment analyses also revealed significant upregulation of pathways specific of the hematopoietic system and the nervous system after downregulation of SA2 in MCF10A cells (Fig. S2), again supporting a role of this variant in establishing and/or maintaining cell identity.

**Figure 2.**
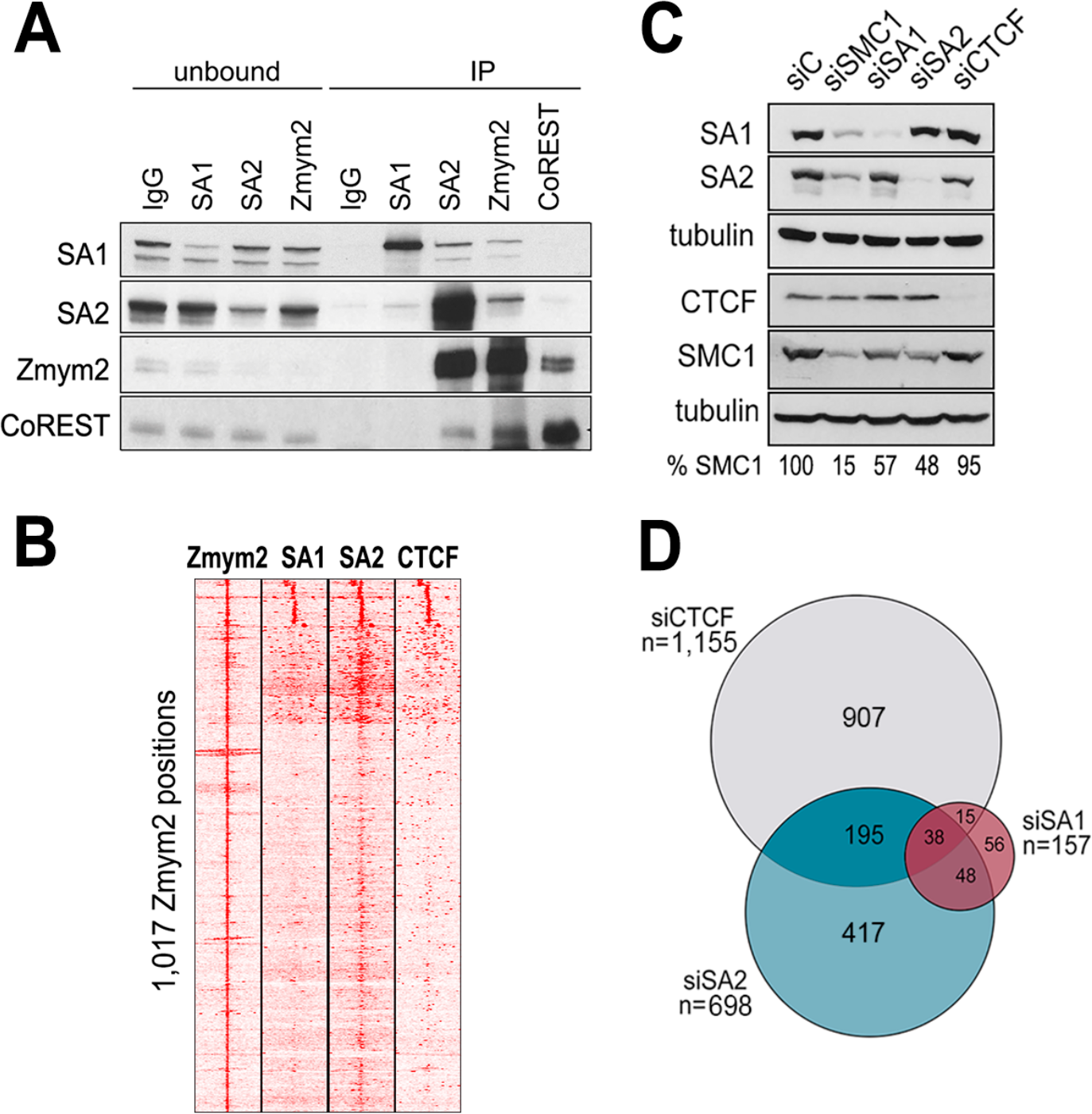
Specific interaction of cohesin-SA2 with transcriptional regulators. (A) Immunoblot analyses of unbound and immunoprecipitated (IP) proteins obtained from HeLa nuclear extracts with the indicated antibodies. (B) Distribution of SA1, SA2 and CTCF around Zmym2 positions (± 5 kb) defined by ChIP-seq in MCF10A cells. (C) Levels of cohesin and CTCF after siRNA transfection in MCF10A cells (siC, mock transfection). (D) Venn diagram showing the overlap between Differentially Expressed Genes (DEGs) after downregulation of SA1, SA2 or CTCF compared to mock transfected cells. To obtain DEGs, FDR<0.05, log_2_fold change<-0.5 or >0.5 and fpkm>3 in at least one of the two conditions compared was required.

Among genes deregulated in siSA2-treated cells there were several members of the S100 family of calcium binding proteins which are located in a 300-kb long gene cluster in chromosome 1 (Table S3 and Fig. S3A). This region contains strong common cohesin peaks at domain boundaries defined by RNApol II ChIA-PET data in mammary cells (*13*) as well as less prominent cohesin-SA2-only binding sites at the promoters of the deregulated genes (Fig. 3A, and S3B). We selected the locus for further analysis and first asked how depletion of the different cohesin subunits affected cohesin binding. The presence of cohesin in common positions (labelled c1 to c4 in Fig. 3A-B) did not decrease in response to SA2 depletion, while both SMC1 and SA2 were significantly reduced in all tested SA2-only positions (labelled o1-o5 in Fig. 3A-B). These results indicate first, that SA1 was unable to replace SA2 in SA2-only positions, and second, that reduced levels of cohesin-SA2 might be irrelevant for the cohesin-CTCF related functions exerted from common positions. Indeed, a 4C experiment using a cohesin c2 position as viewpoint confirmed this hypothesis. Reduced CTCF, but not reduced SA2 levels, impaired the insulation function of such position, as shown by new interactions beyond the domain limit (Fig. S4A). Moreover, after downregulation of SA1, the binding of the remaining cohesin-SA1 to common positions was maintained while binding of cohesin-SA2 decreased significantly (Fig. S4B). As expected, depletion of CTCF reduced the binding of the two cohesin variants (Fig. S4B). The resistance of cohesin-SA1 to be removed from these cohesin-SA1/2-CTCF-binding sites could explain the apparent maintenance of Topologically Associated Domains (TADs) after acute depletion of Rad21 (*14*, *15*) and would agree with a lower turnover of cohesin at those sites. Consistent with this possibility, cohesin complexes at common positions were less likely to associate with Wapl, a factor that dissociates cohesin from chromatin (*16*), compared to cohesin at SA2-only positions (Fig. 3C). We also found that the distribution of cohesin (SMC1) ChIP-seq reads was sharp and narrow around common and cohesin-SA1-only positions in all three cell lines analysed, but much broader around cohesin-SA2-only positions, consistent with the hypothesis of a more dynamically located cohesin in the latter (Fig. 3D). It has been recently proposed that TADs are the result of chromatin bound cohesin complexes acting as loop extruding machines that get stalled at CTCF boundaries (*17*, *18*). We wondered if the stability of cohesin in common positions could be related with the accumulation of more than one cohesin ring at a given CTCF bound site at the base of a chromatin loop. Indeed, Re-ChIP experiments with SA1 and SA2 antibodies revealed that at least two independent cohesin rings can coexist in the same genomic position in the same cell (Fig. 3E). We speculate that higher affinity of SA1 for CTCF and/or lower affinity for Wapl could contribute to the preferential localization of cohesin-SA1 at CTCF sites, which afterwards would stabilize the binding of a newly loaded cohesin-SA2 at the same site (Fig. 3F). This hypothesis concurs with the decrease in SA2 binding at common positions in the absence of cohesin-SA1 (Fig. S4B) as well as our previous observation of an increased number of cohesin positions with lower cohesin occupancy and lower CTCF overlap in SA1 knock out mouse cells (*19*). Our analyses have uncovered a population of cohesin that is present in most cells and is resistant to depletion due to reduced association with Wapl and/or increased stacking of cohesin rings. Cohesin-SA1 together with CTCF have a predominant role in the establishment of this population, which likely drives TAD formation.

**Figure 3.**
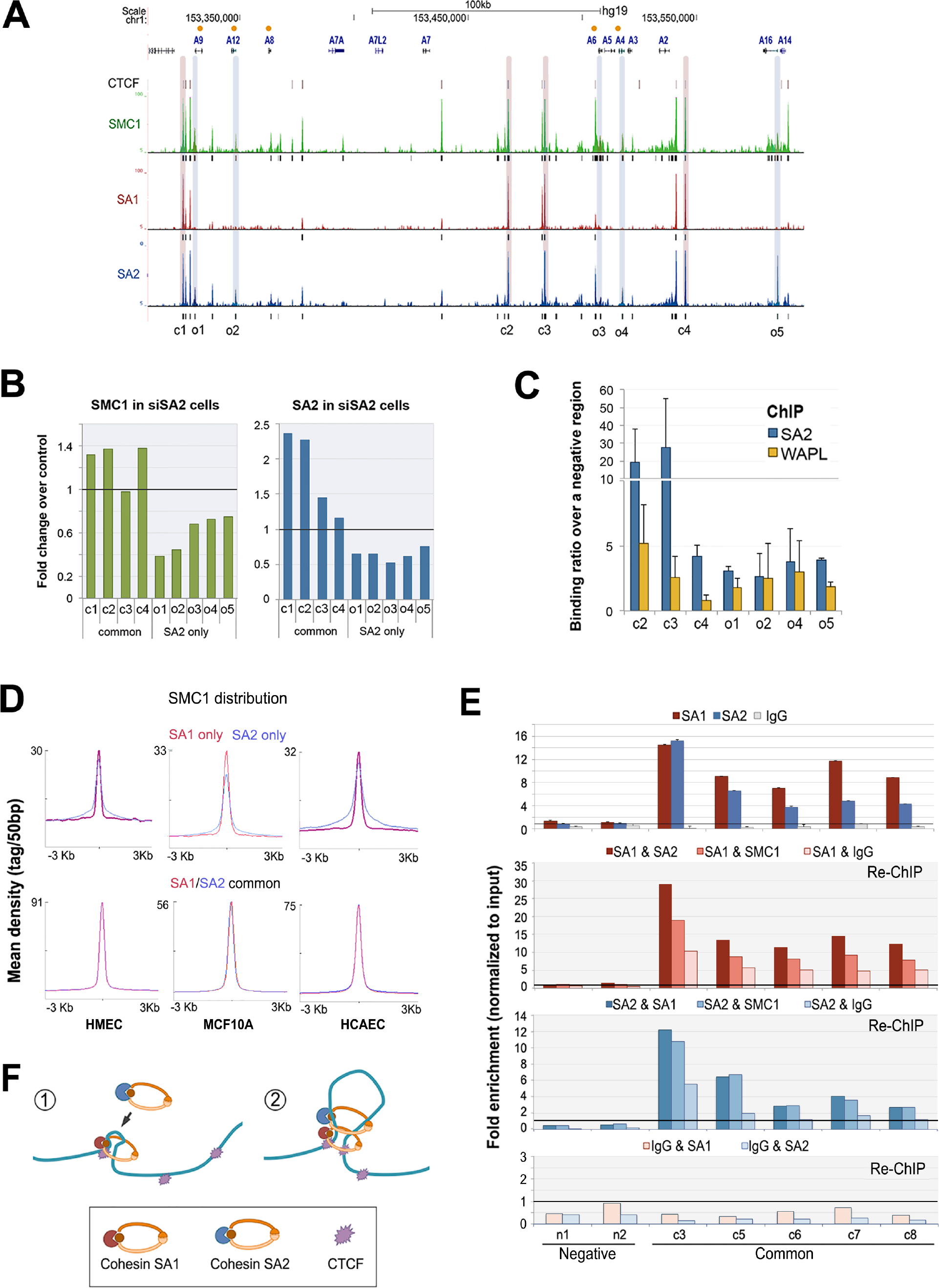
Different behaviour of cohesin in common and SA2-only positions. (A) UCSC browser image of the S100A gene cluster showing genes (orange dots indicate those deregulated in siSA2), CTCF peaks, and genomic distribution of SMC1, SA1 and SA2 in MCF10A cells (aligned reads and called peaks below). Positions corresponding to common (c) and SA2-only (o) cohesin binding sites are shadowed in red and blue, respectively. (B) ChIP-qPCR analyses of cohesin remaining in “c” and “o” positions after downregulation of SA2. SMC1 (green bars) and SA2 (blue bars) ChIP results are represented as fold change of the siSA2 condition over mock transfected condition. (C) ChIP-qPCR analyses of SA2 and WAPL binding to the indicated cohesin positions. Each bar represents the mean of at least three independent experiments performed in triplicates; error bar=SD. (D) Cohesin SMC1 distribution in SA1-only, SA2-only and common positions in HMECs. MCF10A and HCAECs. Maximum mean tag density is indicated on the Y axis. (E) The simultaneous presence of several cohesin complexes at a given position within the same cell was assayed by Re-ChIP. Upper panel, binding of SA1 and SA2 in five common positions (c) as well as two negative regions (n) upon the first immunoprecipitation. IgG ChIP is also shown. Chromatin eluted from the first immunoprecipitation with SA1 or SA2 was incubated with SA2 and SA1 antibodies, respectively, as well as SMC1 and IgG as positive and negative controls. Lower panel, re-ChIP of chromatin eluted from IgG beads with SA1 and SA2. (F) Model for the leading role of cohesin-SA1 in cohesin stacking at the base of a chromatin loop stabilized by converging CTCF bound motifs.

To test this hypothesis, we carried out Hi-C experiments in MCF10A cells depleted from SA1 or SA2 (Fig. S5 and Table S6). Overall, the resulting Hi-C contact maps were similar in all three conditions (Fig. 4A). Identity of active (A) and repressive (B) compartments (*20*) was mostly preserved (Fig. 4B) and the number of TADs was unchanged (Fig. 4C). Consistent with a more prominent role of SA1 in the stabilization of TAD boundaries together with CTCF, TAD border strength was reduced in SA1 depleted cells (Fig. 4D), although this change did not reach statistical significance. The fact that SA2 is also present in common positions, and the resistance of cohesin-SA1 to be removed from them after siRNA treatment (Fig. 3D) likely contribute to dampen the effect of SA1 downregulation. TAD conservation decreased significantly after SA2 depletion (Fig. 4E), which suggests that the formation of a subset of TADs or sub-TADs might occur in a CTCF independent manner and instead depend on the interaction of cohesin-SA2 with different transcriptional regulators. This idea agrees with recent data showing that around 20% of TAD borders are maintained after acute elimination of CTCF in mouse ES cells (*21*). Analysis of the genomic interactions as a function of the genomic distance further evidenced specific contributions of the two cohesin variants to chromatin architecture (Fig. 4F). Loss of SA2 increased mid-range contacts (0.1 -1.3 Mb) that most likely correspond to intra-TAD interactions while loss of cohesin-SA1 increased long-range (>1.4 Mb) inter-TAD interactions. These distinct effects were also evident in the matrices representing differential Hi-C interactions of each condition compared to control cells (Fig. 4G and Fig. S6). Moreover, the increased contacts detected after SA2 depletion occurred in the A compartment and barely affected the B compartment. In contrast, downregulation of SA1 resulted in an overall increase of interactions within B compartment while it decreased interactions within A compartment (Fig. 4H). The specific enrichment of cohesin-SA1 only positions in A/B borders (Fig. 4I) prompt us to speculate that cohesin-SA1 may play a unique role in modulating A/B compartment identity, in addition to or as a consequence of its major role in TAD border definition. Cohesin-SA2 would be more critical for intra-TAD contacts, and its loss would lead to new stochastic interactions between enhancers and nearby promoters, increasing overall chromatin contacts and altering gene expression (see model in Fig. 4J).

**Figure 4.**
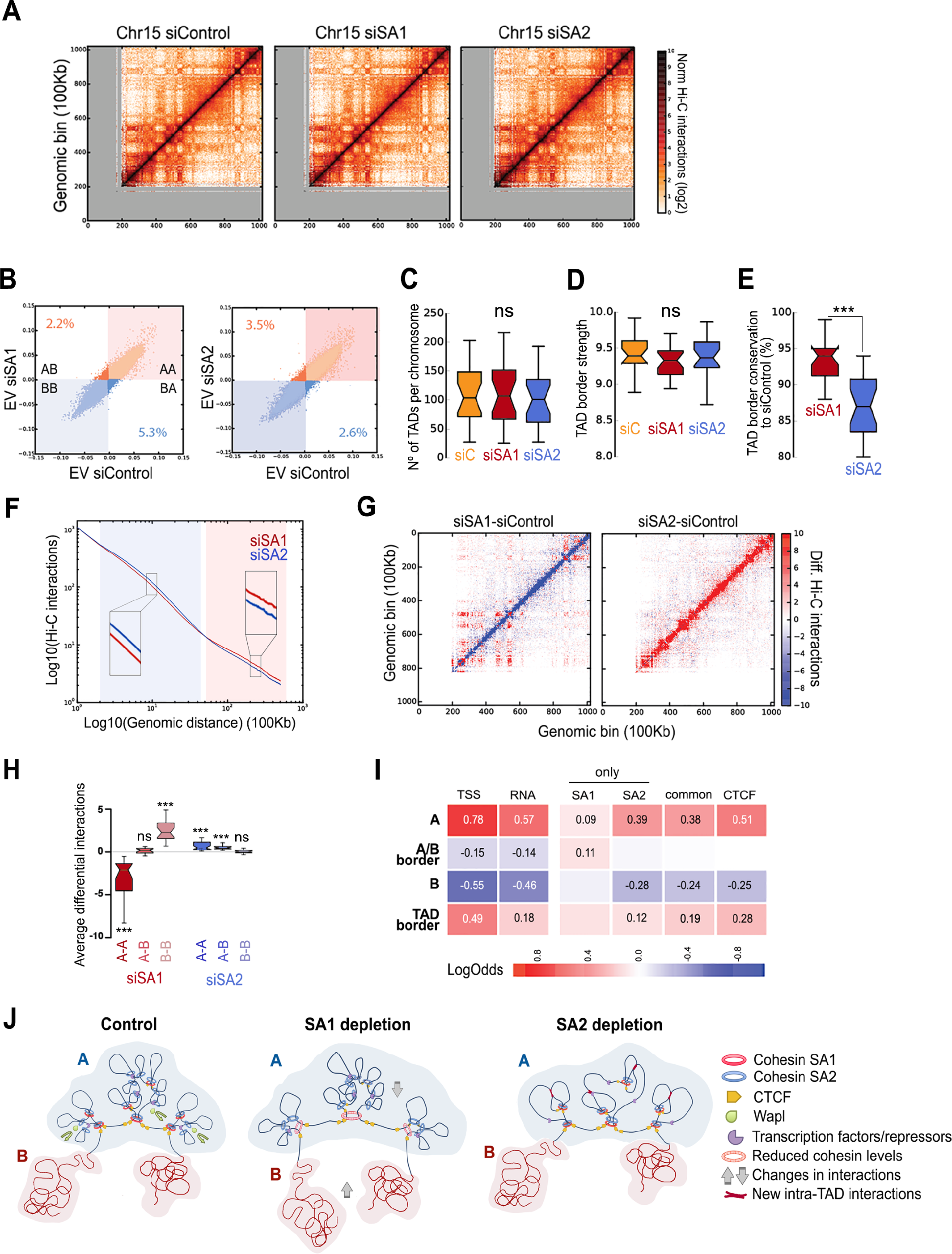
Distinct contribution of cohesin-SA1 and cohesin-SA2 to genome architecture. (A) Hi-C normalized interaction maps for chromosome 15 at 100Kb resolution in MCF10A cells mock transfected (siControl), or treated with siSA1 or siSA2. (B) Scatter plot of Eigen vectors of the intra-chromosomal interaction matrices indicated. Numbers within the plot are the % of 100Kb bins that change compartment. (C) Box plot of number of TADs per chromosome and (D) TAD border strength in the three conditions indicated. (E) TAD border conservation in siSA1 and siSA2 with respect to control. (F) Chromosome Hi-C interactions as a function of genomic distance averaged across the genome for maximum distances of 50 Mb. Background colours highlight the intervals where higher interactions are detected for siSA1 (red) and siSA2 (blue). (G) Chromosome 15 differential interactions matrix comparing siControl to siSA1 or siSA2 in and (H) genome-wide box-plot of chromosome differential interactions separated by A-A, A-B and B-B contacts; ***p-value<0.001. Separate analyses of the two replicates are shown Fig. S6. (I) Enrichment/depletion (Log Odds) of SA1-only, SA2-only and common cohesin binding sites in AB compartments, A/B borders and TAD borders. Squares with numbers are significant (Fisher exact test, p-values < 0.005). (J) According to the "loop extrusion" model, once cohesin-SA1 is loaded onto chromatin, it extrudes the DNA fibre until it encounters two CTCF molecules disposed in the convergent orientation. The cohesin ring will stop at that position and stabilize the resulting loop. Additional cohesin complexes, including those containing SA2, can accumulate at those positions, reinforcing the loop stability and generating insulation regions that will define TAD or subTAD boundaries. Cohesin-SA2 will be retained more easily by transcription factors other than CTCF and will stabilize loops within TADs that support interactions between enhancers and their target genes. These loops are more transient since Wapl promotes dissociation of cohesin. Depletion of SA1 leaves a significant fraction of cohesin-SA1 at TAD borders. Even though TAD border strength is reduced, no major impact on TAD identity is observed. However, genomic compartmentalization is altered by increasing the interactions within B compartment at the expense of interactions within A compartment. Cohesin-SA2 depletion, in contrast, disorganizes intraTAD chromatin loops, increasing local interactions many of them probably aberrant (“new intraTAD interaction”) and thereby alters gene expression.

Recent studies depleting cohesin or CTCF in different cellular systems have led to the conclusion that TADs and compartments arise independently (*21*, *22*). TADs would depend on cohesin and CTCF, whereas genomic compartmentalization would rely mostly on epigenetic features regardless of chromatin contacts. While TAD boundaries are largely invariant across cell types, the specific interactions within TADs may not (*23*, *24*). Moreover, single-cell HiC experiments imply a certain degree of stochasticity in the TAD boundary definition among cells in the population (*25*, *26*). In this study, we show for the first time that the two variant cohesin complexes present in somatic cells, cohesin-SA1 and cohesin-SA2, make distinct contributions to higher order chromatin organization. Taken together the features of cohesin-SA2 only positions in the genome, the physical interaction of SA2 with transcriptional regulators such as CoREST complex and intra-TAD contact alteration resulting from downregulation of SA2, we propose that cohesin-SA2 regulates tissue-specific gene expression by modulating the spatial organization of enhancers and promoters within TADs. Cohesin-SA1, in contrast, would drive stacking of cohesin rings at CTCF-bound sites to stabilize TAD boundaries. Our results further indicate that cohesin-SA1 cannot fully compensate the loss of SA2. While its presence likely allows *STAG2* mutant cancer cells to survive, it is insufficient to properly regulate gene expression.

## Materials and Methods

### Cell lines

All the human primary cell lines in this study were purchased from Lonza and cultured according to the manufacturer’s recommendations. NHA (Normal Human Astrocytes, CC- 2565) were grown in ABM basal medium (CC-3187) supplemented with AGM Bulletkit (CC-4123); SKMC (Skeletal Muscle Cells, CC-2561) were cultured in SkBM basal medium (CC-3161) supplemented with SkGM Bulletkit (CC-4139); NHBE (Normal Human Bronchial Epithelial Cells, CC-2540) were cultured in BEBM basal medium (CC- 3171) supplemented with BEGM Bulletkit (CC-4175); HCAEC (Coronary Artery Endothelial Cells, CC-2585) were grown in EBM2 basal medium (CC-3156) supplemented EGM2-MV Bulletkit (CC-4147); NHEK (Normal Human Epidermal Keratinocytes, #00192627) were grown in KBM-Gold basal medium (#00192151) supplemented with KGM-Gold Bulletkit (#00192060). HMEC (Normal Mammary Epithelial Cells, CC-2551) were cultured in MEBM basal medium (CC-3171) supplemented with MEGM Bulletkit (CC-3150). NHOst (Normal Human Osteoblasts) were grown in OBM basal medium (CC- 3208) supplemented with OGM Bulletkit (CC-3207). PrEC (Prostate Epithelial Cells, CC- 2555) were cultured with PrEBM basal medium (CC-3165) supplemented with PrEGM Bulletkit (CC-3166). HUVEC (Human Umbilical Vein Endothelial Cells, CC-2517) were grown in EBM basal medium (CC-3121) supplemented with EGM Bulletkit (CC-3124). MCF10A cells were kindly provided by M. Quintela, (CNIO, Madrid) and grown in DMEM/F12 (#31330038, Thermofisher) supplemented with 20ng/ml of EGF, 0.5mg/ml hydrocortisone, 100ng/ml of cholera toxin, 10mg/ml of insulin and 5% of horse serum.

### Antibodies

A rabbit polyclonal antibody recognizing human Wapl was generated using a recombinant C-terminal fragment of the protein (352 amino acids long) obtained by PCR amplification of full length hWapl cDNA [a gift from T. Hirano (RIKEN, Japan)]. Additional custom made antibodies have been previously described: SA1, SA2 and SMC1 (***27)***, RAD21 (***28)***, Zmym2 [a generous gift from H. Yu (UT Southwestern, US)] (***29)***. Commercial antibodies were CTCF (07-729, Millipore), CoREST (07-455, Millipore), tubulin (DM1A, Sigma), histone H3 (Abcam AB1791).

### Immunoblotting

To analyze protein levels in the different cell lines, cells were collected by trypsinization, counted, washed once in cold PBS, resupended in SDS-PAGE loading buffer at 10^7^ cells/ml, sonicated and boiled. Equal volumes were separated by SDS-PAGE and analyzed by Western blotting. Chromatin fractionation was performed as described (***30)*** and fractions were run on SDS gels alongside increasing amounts of recombinant proteins corresponding to C-terminal fragments of human SA1 and SA2 to estimate the amount of each variant subunit in the chromatin fractions, as in (***27)***.

### Immunoprecipitation and LC-MS/MS Analysis

Whole cell extracts from MCF10A cells were prepared by lysis on ice for 30 min in TBS supplemented with 0.5% NP-40, 0.5mM DTT, 0.1mM PMSF and 1X complete protease inhibitor cocktail (Roche) followed by sonication. NaCl was added to 0.3M and the extract rotated for 30 min at 4ºC. After centrifugation, the soluble fraction was recovered and diluted to bring the extract back to 0.1M NaCl and 10% glycerol was added. HeLa nuclear extracts were prepared from isolated nuclei as previously described ***(7)***. Antibodies were cross-linked to protein A Pureproteome magnetic beads (Millipore) at 10 mg/ml (SA1, SA2 and IgG as control). Extracts were rotated overnight at 4ºC with antibody-beads. The beads were washed 3 times with 20 vol of buffer containing 10mM K-Hepes pH 7.7, 0.1M KCl, 2mM MgCl_2_, 0.1mM CaCl_2_, 5mM EGTA and 3 times with the lysis buffer. They were eluted in two consecutive steps in 2 vol of elution buffer (8M urea, 100mM Tris-HCl pH by shaking for 10 min. Samples were digested by standard Filter Aided Sample Preparation (FASP) **(*31*)**. Proteins were reduced with 10mM DTT, alkylated with 50mM IAA for 20 min in the dark and digested with 1:50 Lys-C (Wako) for 4 h. Samples were diluted in 50mM ammonium bicarbonate and digested with 1:100 Trypsin (Promega) overnight at 37ºC. Resulting peptides were desalted using a Sep-Pak C18 cartridge for SPE (Waters Corp.), vacuum-dried and resuspended in 0.5% FA. Immunoprecipitates from HeLa nuclear extracts were analyzed by RP chromatography using a nanoLC Ultra system (Eskigent) directly coupled with the Impact instrument (Bruker Daltonics) equipped with a CaptiveSpray nanoelectrospray ion source supplemented with a CaptiveSpray nanoBooster operated at 0.2 bar/minute with isopropanol as dopant. Two experiments and two technical replicates (runs) of each were performed. Immunoprecipitates from MCF10A extracts were analysed using a nanoLC Ultra system (Eksigent, Dublin, CA) coupled with a LTQ-Orbitrap Velos instrument (Thermo) via nanoESI (ProxeonBiosystem, Waltham, MA). A single immunoprecipitation experiment and two technical replicates were performed in this case. Raw data were analysed using MaxQuant (***32***) 1.5.3.30 with Andromeda (***33***) as the search engine against UniProtKB/Swiss-Prot (20,584 sequences). Peptides were filtered at 1% FDR. For protein assessment (FDR <1%), at least one unique peptide was required for both identification and quantification. Other parameters were set as default. The resulting “proteingroup.txt” file was loaded in Perseus (***34***)(v1.5.1.6). Missing values were imputed from a normal distribution. A two-sample Student’s T-Test (one side) was used corrected for multiple testing using a permutation-based approach. For HeLa, significant enrichment was set at FDR<0.05, S0=0.1 for SA1_IgG and FDR<0.01, S0=0.8 for SA2_IgG (in HeLa cells cohesin-SA2 is much more abundant than cohesin-SA1). In MCF10A, FDR<0.05, S0=0.1 was used. A detailed protocol for LC-MS/MS Analysis can be provided upon request.

### siRNA

MCF10A cells were transfected with 50 nM onTARGETplus SMARTpool siRNAs (Dharmacon L-010638, L-021351, L-006833 and L-020165 for SA1, SA2, SMC1 and CTCF, respectively) using DharmaFECT reagent 1. Transfection efficiency was first estimated by qRT-PCR 24 h after transfection, and typically reached more than 90% downregulation. Cells were taken at 72h and protein levels assessed by immunoblot.

### ChIP sequencing and analysis

Chromatin immunoprecipitation (ChIP) was performed as described **(*35*)**, with some modifications. Confluent cells were cross-linked with 1% formaldehyde added to the media for 15 minutes at RT. After quenching with 0.125M Glycine, fixed cells were washed twice with PBS containing 1uM PMSF and protease inhibitors, pelleted and lysed in lysis buffer (1%SDS, 10mM EDTA, 50mM Tris-HCl pH 8.1) at 2x10^7^ cells/ml. 10^7^ cells equivalent to 40-50 mg of chromatin were used per immunoprecipitation reaction with xx mg of antibody. Sonication was performed with a Covaris system (shearing time 30 min, 20% duty cycle, intensity 6, 200 cycles per burst and 30 s per cycle) in a minimum volume of 2 ml. From 6 to 8 ng of immunoprecipitated chromatin (as quantitated by fluorometry) were electrophoresed on an agarose gel and independent sample-specific fractions of 100–200 bp were taken. Adapter-ligated library was completed by limited-cycle PCR with Illumina PE primers (13 cycles). DNA libraries were applied to an Illumina flow cell for cluster generation and sequenced on the Illumina Genome Analyzer IIx (GAIIx). Image analysis was performed with Illumina Real Time Analysis software (RTA1.8).

Alignment of 50-bp long sequences to the reference genome (GRCh37/hg19, February 2009) was performed using ‘BWA’ **(*36*)** under default settings. Duplicates were removed using Picardtools (version 1.60) and peak calling was carried out using MACS2 (version 2.1.1.20160309) setting a q value (FDR) to 0.05 or 0.01 (SMC1,SA1, SA2 in HMEC) and using the ‘--extsize’ argument with the values obtained in the ‘macs2 predictd’ step (***37)***. All comparisons used the input tracks as "control", and each one of the datasets as "treatment".

Common, SA1-only and SA2-only positions were defined using BEDtools v2.26 with a minimum of 1 nt overlap. Common positions were defined in two steps: 1) overlap between SMC1 and SA1 bed files was performed using ‘-wa -wb’ argument and the positions obtained were concatenated and sorted using ‘cat’ and ‘sort -k1,1 -k2,2n’ commands. The output was merged using ‘bedtools merge’ function and considered as one dataset; 2) this was overlapped with the SA2 dataset as above. SA1-only and SA2-only positions are those where SA1 or SA2 do not overlap among each other.

Mean read density profiles and read density heatmaps for different chromatin binding proteins were generated with seqMINER v1.3.3e using BAM files of processed reads and plotting them around peak summits of SA1 or SA2 only or common positions.

For Motif discovery analysis, whole sequences of cohesin positions were extracted and used for motif enrichment analysis using MEME-ChIP from MEME **(*38*)**. Default parameters were used except for the following ones: -ccut 0, -meme-mod anr, -meme-minw: 6, -meme-maxw: 50, -nmeme: 600, -meme-nmotifs: 10, -meme-maxsize: 200,000.

Enrichment of cohesin positions (SA1 and SA2-only and common) at HMEC chromatin states was defined using ‘intersect’ function from BEDtools utilities (v2.26) with a minimum of 1nt overlap. The analysis was performed making sure that one position does not belong to two different chromatin states.

### ChIP-qPCR and ReChIP

ChIP-qPCR on immunoprecipitated chromatin was performed using the SYBR Green PCR Master Mix and an ABI Prism® 7900HT instrument (Applied Biosystems®). Primers were designed using OligoPerfect Designer™ (Invitrogen) and reactions were performed in triplicate. Quantifications were normalized to endogenous GAPDH, using the ΔΔCt method. Chromosome coordinates of the validated peaks and the corresponding primers are listed in Supplementary Table S6. The relative amount of each amplified fragment was normalized with respect to the amplification obtained from input DNA and represented as indicated in the corresponding figure legends.

ReChIP experiment was performed with the Re-ChIP-IT kit (#53016, Active Motif) according to the manufacturer’s protocol. Briefly, MCF10A cells were fixed, lysed and sonicated as described in the ChIP protocol. 50 mg of chromatin were incubated with 20 μg of the first antibody (SA1, SA2 or IgG) in presence of magnetic beads, washed, eluted and further incubated with 5 mg of the second antibody (SA1, SA2, SMC1 or IgG). Eluted chromatin was analyzed by quantitative PCR as described above (primers in Table S6).

### Quantitative RT-PCR and RNA-sequencing

Total RNA was extracted using the RNeasy Mini Kit (Qiagen) and cDNAs were prepared according to the manufacturer’s instructions using the Superscript II reverse transcriptase (Invitrogen). qRT-PCR analysis was performed using the SYBR Green PCR Master Mix and an ABI Prism® 7900HT instrument (Applied Biosystems®). Primers (Table S6) were designed using OligoPerfect Designer™ (Invitrogen) and reactions were performed in triplicate. Quantifications were normalized to endogenous GAPDH, using the ΔΔCt method.

For RNA-seq libraries (three replicates for condition), RNA was extracted as described and treated with DNaseI (Ambion). polyA+RNA was purified with the Dynabeads mRNA purification kit (Invitrogen), randomly fragmented and converted to double stranded cDNA and processed through subsequent enzymatic treatments of end-repair, dA-tailing, and ligation to adapters as in Illumina’s "TruSeq RNA Sample Preparation Guide" (Part # 15008136 Rev. A). Adapter-ligated library was completed by limited-cycle PCR with Illumina PE primers (8 cycles). The resulting purified cDNA library was applied to an Illumina flow cell for cluster generation (TruSeq cluster generation kit v5) and sequenced on the Genome Analyzer IIx with SBS TruSeq v5 reagents by following manufacturer’s protocols. Fastq files with 50-nt single-end sequenced reads were quality-checked with FastQC (S. Andrews, http://www.bioinformatics.babraham.ac.uk/projects/fastqc/) and aligned to the human genome (GRCh37/hg19) with Nextpresso (http://bioinfo.cnio.es/nextpresso/) executing TopHat-2.0.0 using Bowtie 0.12.7 and Samtools 0.1.16 allowing two mismatches and five multi-hits. Transcript assembly, estimation of their abundances and differential expression were calculated with Cufflinks 1.3.0 using the mouse genome annotation data set GRCm37.v65 from Ensembl. To account for multiple hypotheses testing, the estimated significance level (*p* value) was adjusted using Benjamini-Hochberg False Discovery Rate (FDR) correction. For differential expression, FDR<0.05, log2fold change<-0.5 or >0.5 and fpkm>3 in at least one of the two conditions compared was required.

GSEAPreranked was used to perform a gene set enrichment analysis **(*39*)**. We used the RNA-seq gene list ranked by statistic, setting ‘gene set’ as the permutation method and we run it with 1000 permutations.

### 4C-seq

Preparation of 4C-seq samples was performed as described **(*40*)** with some modifications. Cells were cross-linked with 2% formaldehyde for 10 min at RT on a rocking platform and quenched by addition of glycine to 0.125M and incubation for 5 min at RT. After washing twice with cold PBS, cells were harvested in cold PBS containing 100 mM PMSF and protease inhibitor cocktail. Cells were recovered by centrifugation at 1,300 rpm for 8 min at 4°C and resuspended in lysis buffer (50-mM Tris pH 7.5, 150-mM NaCl, 5-mM EDTA, 0.5% NP-40, 1% TX-100 + protease inhibitors cocktail) at a density of 20x10^6^ cells/ml. After 20 minutes at 4°C for, lysis was assessed by methyl green-pyronin staining. DpnII and CviAII restriction enzymes were used as first and second 4-cutter, respectively. 4C- seq libraries were amplified using long primers with 20 bp homology to the bait sequence and Illumina paired-end adapter flanks. Primer sequences at the viewpoint site are as follows: FW 5′-CCCCCAAGGTCATAACGATC-3′, and Rev 5′-ATTGTCCCAGTGATGGGCAG-3′. 4C-seq data analysis and normalization was performed using HTS station on the hg19 human genome assembly.

### Hi-C

MCF10A cells were arrested in G1 by means of high confluency culture (150,000 cells per cm^2^). Hi-C was performed as described ***(41)*** using MboI enzyme. Two library replicates per condition were sequenced (>200 million reads each, see Table S6). Data were processed using TADbit ***(42)*** for read quality control, read mapping, interaction detection, interaction filtering, and matrix normalization. First, the reads were checked using an implemented FastQC protocol in TADbit. This allowed discarding problematic samples and detect systematic artefacts. Then, we used a fragment-based strategy in TADbit for mapping the remaining reads to the reference human genome (gr38). The mapping strategy resulted in about 80% of reads mapped uniquely to the genome. Next, we filtered non-informative contacts between two reads, including self-circles, dangling-ends, errors, random breaks or duplicates as previously described ***(42)***. The final interaction matrices resulted in 272 to 303 millions of valid interactions per experimental condition (Table S6, Fig. 4A). These valid interactions were then used to generate genome-wide interaction maps at 100 Kb, 40 Kb and 10Kb to segment the genome into the so-called A/B compartments, Topologically Associating Domains (TADs), and produce differential interaction maps.

A/B compartments were calculated using vanilla normalized and decay corrected matrices (41) as implemented in TADbit (42). Briefly, compartments are detected by calculating the first component of a PCA of chromosome-wide matrices and assigning A compartments to genomic bin with positive PCA1 values and high genes density (Fig. 4B). Conversely, B compartments are assigned to genomic bin with negative PCA1 values and low genes density. TADs were identified using 40 Kb resolution vanilla normalized and decay corrected matrices as input to the TAD detection algorithm implemented in TADbit. TAD border localization as well as strength was calculated and used to identify conserved borders and their strength (Fig. 4C-E). A border was considered conserved between siControl and siSA1 or siSA2 experiments if it was localized within +/- 2 two bins in both experiments. Raw matrices normalized by coverage (that is, all three experiments were scaled to have the same number of final valid interactions) at 100Kb resolution were also used for studying Hi-C interactions as function of genomic distance. This genomic decay was obtained per chromosome to a maximum genomic distance of 50Mb and then averaged to obtain a genome-wide curve in siSA1 and siSA2 experiments in (Fig. 4F). The same 100Kb matrices were used to determine differential Hi-C interactions between siControl and siSA1 or siSA2 experiments (Fig. 4G). These differential interactions maps were then used to assess the chromosome average differential interaction as a function of compartment localization (Fig. 4H). Finally, the enrichment or depletion of genes (represented by their TSS), RNA (based on RNA-seq data), and CTCF and cohesin binding sites (SA1-only, SA2 and common) was analyzed by a log odds analysis of observing such features in genomic bins belonging to A, B compartments, A/B borders or TAD borders (Fig. 4I). The log odds distributions were assessed for their distribution being statistically different than zero as for a Fisher exact test (p-value <0.005).

### Data access

ChIP-seq, RNA-seq, 4C-seq and Hi-C data from this study have been submitted to GEO database (GSExxxxxx).

## Acknowledgements

### Acknowledgments

We thank Yasmina Cuartero and Javier Quilez (4D Genome- CRG) for their technical help with the Hi-C experiments, and Nuria Ibarz and Javier Muñoz (CNIO) for proteomic analyses. We are also indebted to Daniel Rico (Newcastle University), Francisco X. Real (CNIO) and Miguel Manzanares (CNIC) for their comments to the manuscript; to members of our group and the group of Juan Méndez (CNIO) for useful discussions; and to Tatsuya Hirano (RIKEN) and Hongtao Yu (UT Southwestern) for reagents. This work has been supported by the Spanish Ministry of Economy and Competitiveness and FEDER funds (grants BFU2013-48481-R and BFU2016-79841-R to AL; BFU2013-47736-P to MAM-R; BES-2014-069166 fellowship to MDK; Centro de Excelencia Severo Ochoa SEV-2015-0510 to CNIO and SEV-2012-0208 to CRG); the European Research Council under the 7th Framework Program (FP7/2010-2015, ERC grant agreement 609989) and the European Union’s Horizon 2020 research and innovation programme (agreement 676556) to MAM-R; CERCA Programme/Generalitat de Catalunya to MAM-R; La Caixa Foundation (PhD fellowship to AK). The authors declare no conflict of interest.

---

**Fig. S1.**
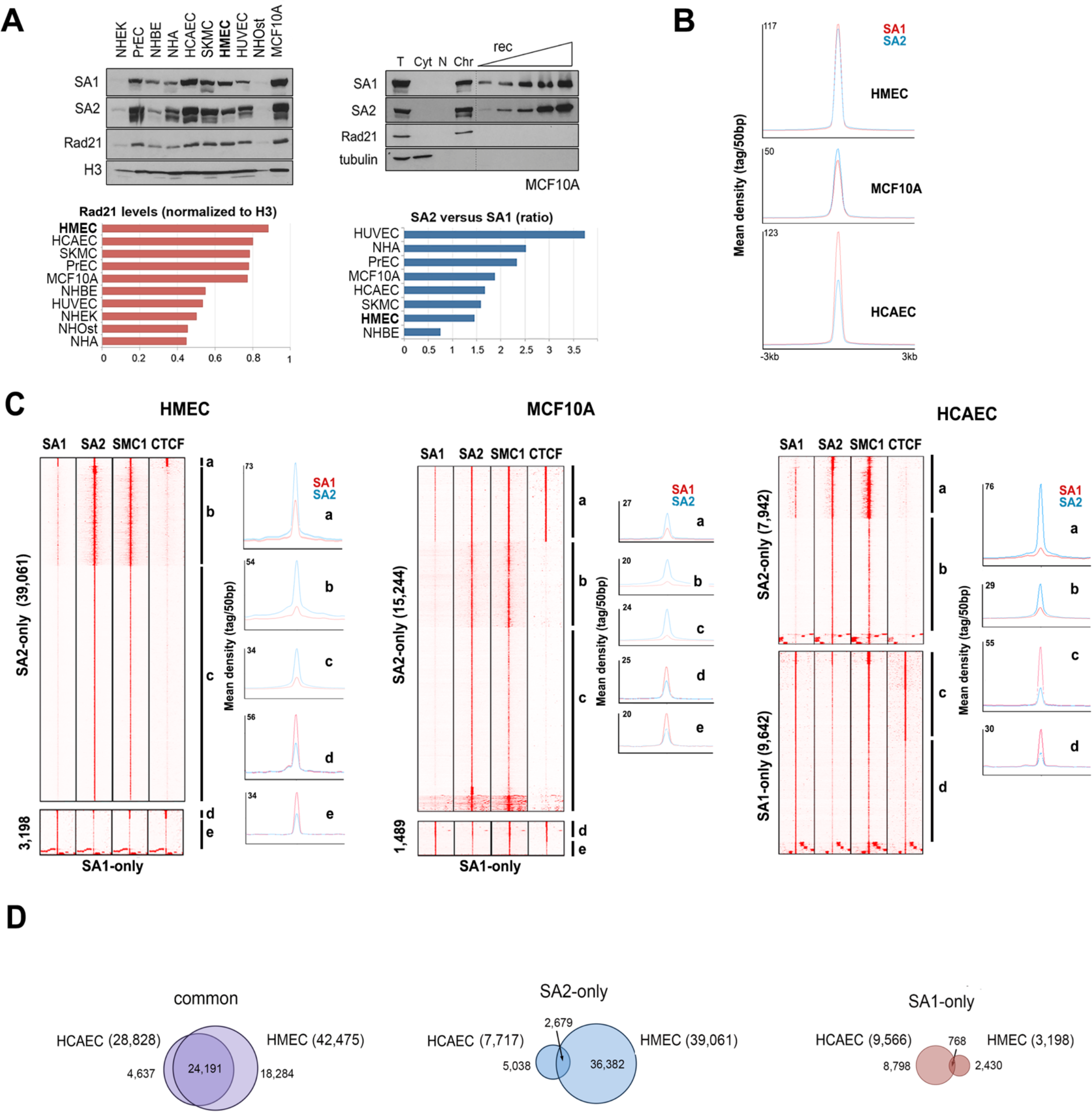
Distinct genome-wide distribution of cohesin-SA1 and cohesin-SA2 in human cells. (A) Top left, cohesin variant abundance in human primary cells (NHEK, Normal Human Epithelial Keratinocytes; PrEC, Prostate Epithelial Cells; NHBE, Normal Human Bronchial Epithelial Cells; NHA, Normal Human Astrocytes; HCAEC, Human Coronary Artery Endothelial Cells; SKMC, Skeletal Muscle Cells; HMEC, Human Mammary Epithelial Cells; HUVEC, Human Umbilical Vein Endothelial Cells; NHOst, Normal Human Osteoblasts) and MCF10A. Top right, known amounts of C-terminal fragments of SA1 and SA2 were run alongside the indicated fractions from MCF10A cells to assess relative abundance of cohesin-SA1 and cohesin-SA2 on chromatin (T, total cell extract; Cyt, cytosolic fraction; N, Soluble Nuclear Fraction; Chr, Chromatin bound protein). This was repeated with all cell lines used above (not shown). Bottom, histograms showing total cohesin levels and SA2:SA1 ratio in the same cell lines. (B) and (C) Average read density plots for SA1 (red) and SA2 (blue) distribution in (B) common positions and (C) defined clusters within SA1-only and SA2-only positions (labelled in lower case) in HMECs, MCF10A cells and HCAECs. Maximum mean tag density is indicated in the Y axis. Identical extent of cohesin enrichment around common positions suggests a similar efficiency of the SA1 and SA2 antibodies. Note that while in most SA2-only positions SA1 levels are minimal, SA1-only positions have a significant number of SA2 reads. (D) Venn diagrams showing overlap of common, SA1-only and SA2-only cohesin binding sites between two cell lines of distinct embryonic origin, HMECs and HCAECs. Common positions are clearly more conserved.

**Fig. S2.**
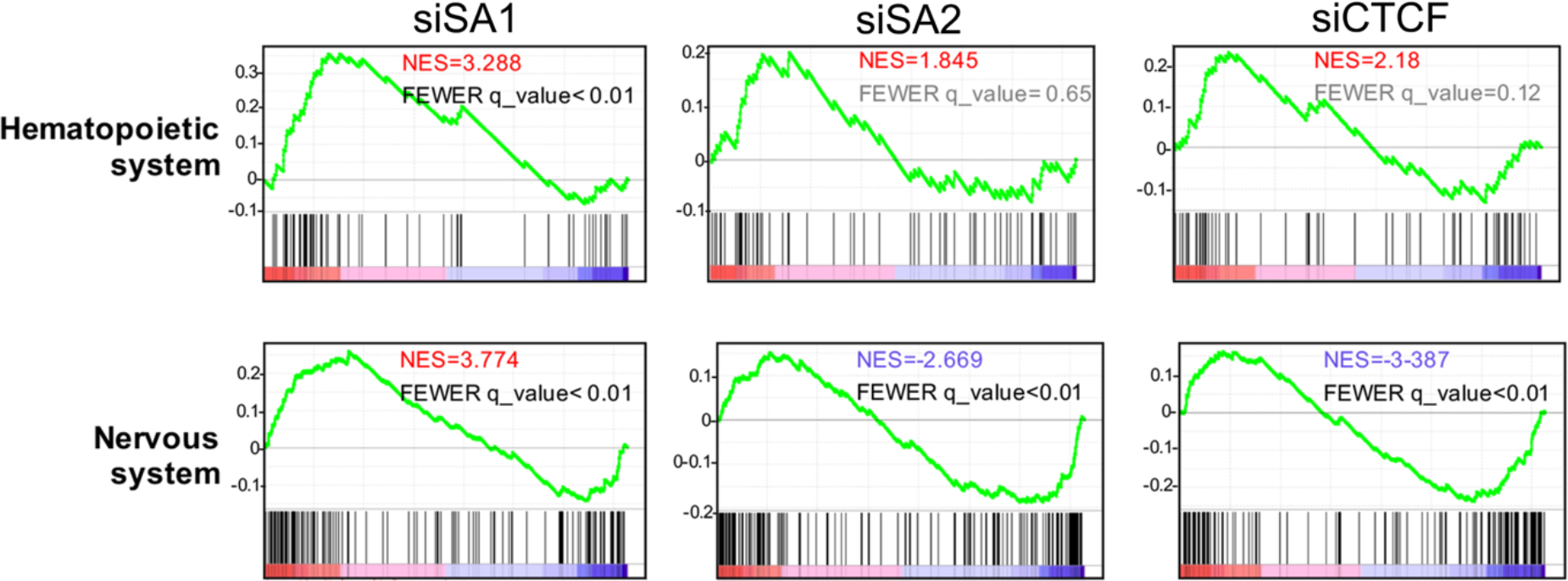
Cohesin-SA2 contributes to maintain cell identity. Enrichment plots for genes within KEGG pathways specific for the hematopoietic system (“Hematopoietic cell lineage”) and the nervous system (“Neuroactive Ligand-Receptor interaction”) after downregulation of SA1, SA2 or CTCF. Both these pathways appear significantly upregulated in the siSA2 condition (NES>0, in red; FWER q-value<0.01). In the other two conditions the gene expression changes are either not significant (FWER q-value in grey) or in the opposite direction (NES<0, in blue). Y axis, enrichment score (ES).

**Fig. S3.**
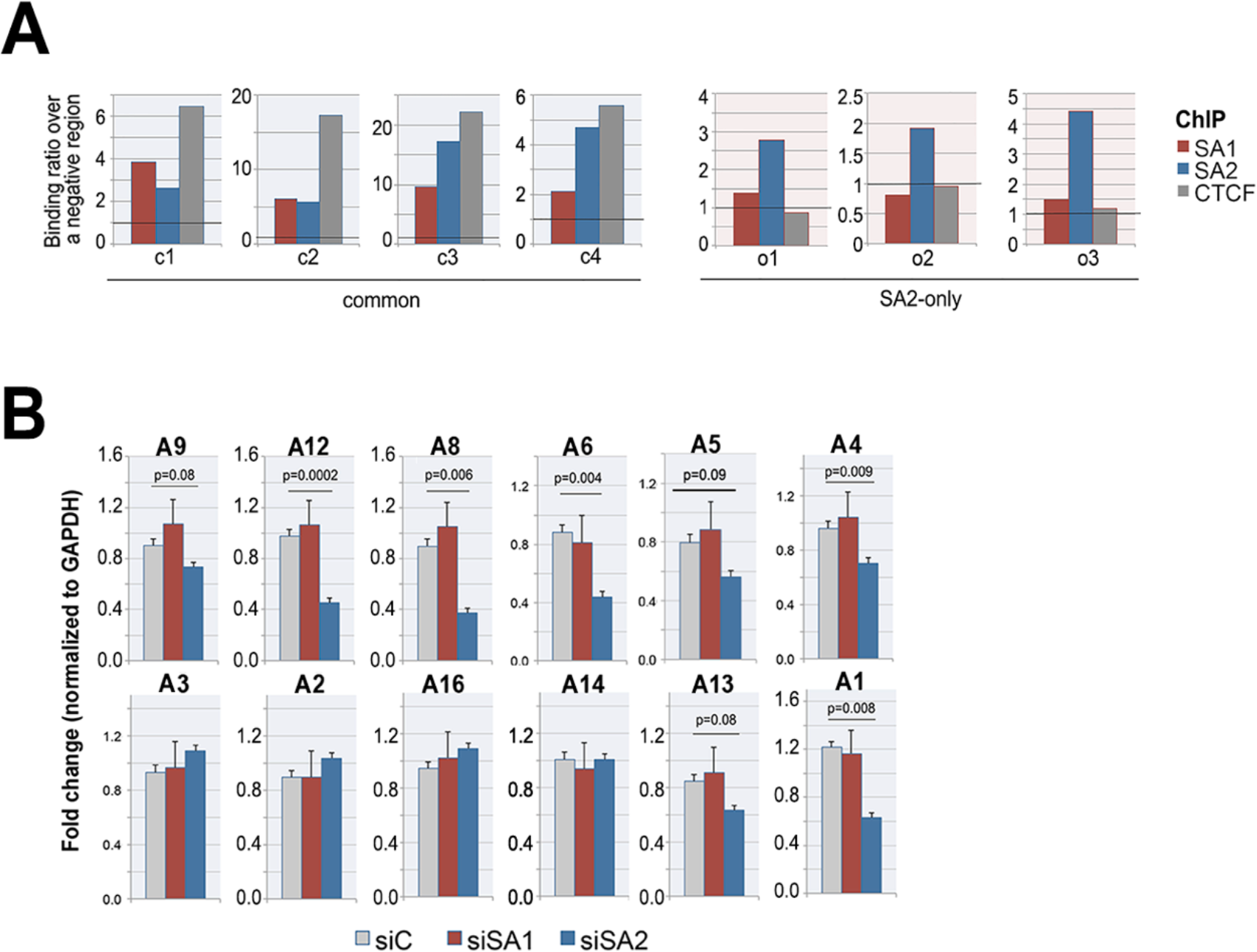
Cohesin distribution around the S100A gene cluster. (A) ChIP-qPCR validation of SA1, SA2 and CTCF binding to the common (c) and SA2- only (o) positions of the S100A locus indicated in Fig. 3A. Binding in each position was calculated as fold increase over the binding to a negative genomic region upon normalization to input. (B) Gene expression levels of S100A genes in control, siSA1 and siSA2 conditions. RNA from three independent samples per condition was used. SA1 depletion did not affect gene expression, while SA2 did (orange dots in Fig. 3A). Student’s T test was used to assess statistical significance.

**Fig. S4.**
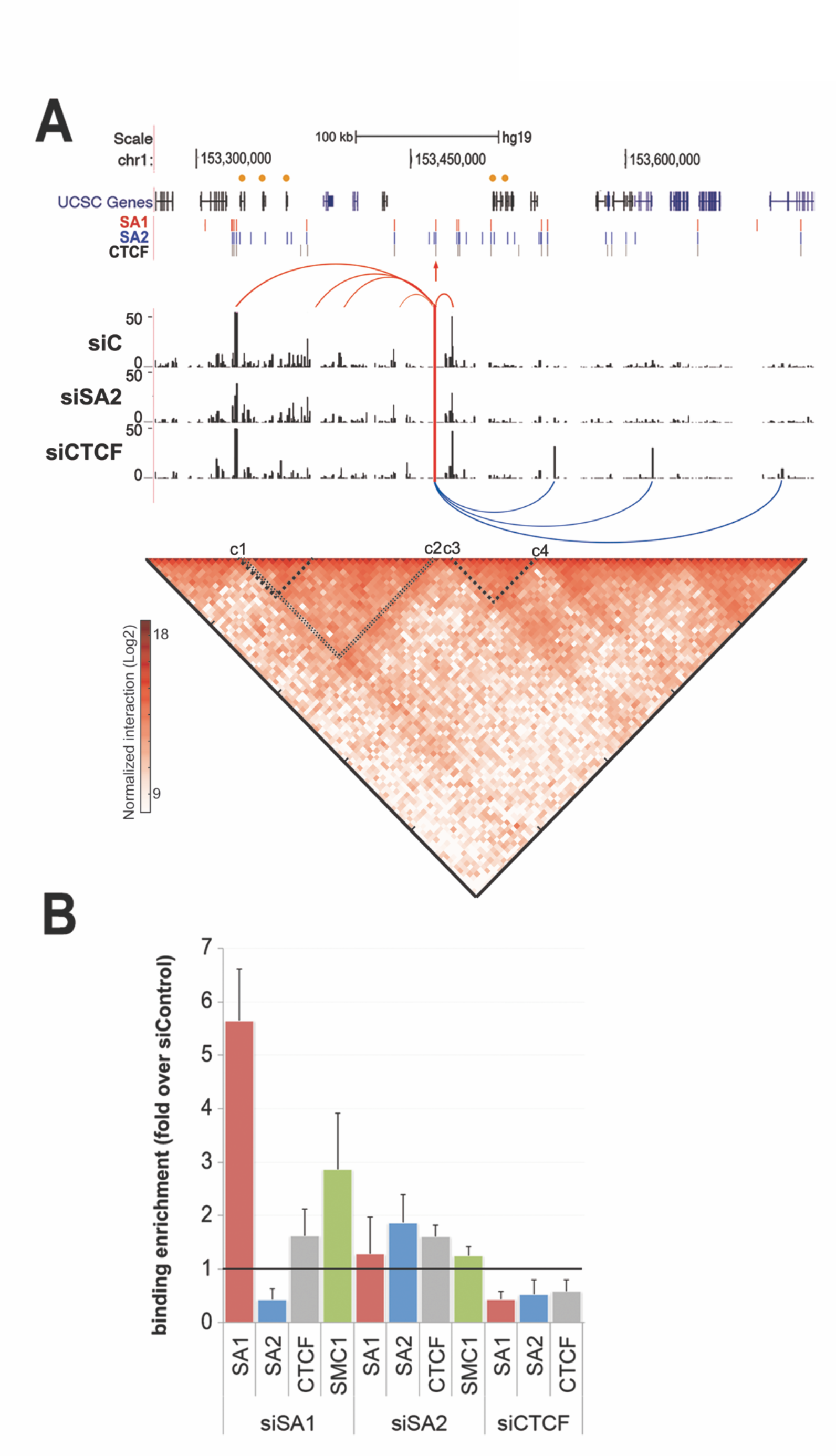
Common cohesin positions are resistant to depletion. (A) A 4C experiment in S100A locus was performed in MCF10A cells with reduced SA2 or CTCF levels. A vertical red line and an arrowhead indicate the viewpoint position. Binding sites of cohesin-SA1, cohesin-SA2 and CTCF identified by ChIP-seq are also shown. Red curves represent interactions present in the mock transfected cells. Blue curves indicate new interactions detected after CTCF depletion. Bottom, Hi-C contact matrix from S100A locus at 5kb resolution in MCF10A cells. The intensity of each pixel represents the normalized number of contacts between a pair of loci. Maximum intensity is indicated. Dotted lines highlight regions with high contact frequency. (B) Changes in cohesin (SA1, SA2, SMC1) and CTCF binding after depletion of SA1, SA2 or CTCF were estimated by ChIP-qPCR. Bars show the average fold change in five common positions (c1 to c5) for each condition compared to mock transfected cells. Strikingly, a clear increase of cohesin-SA1 was observed after downregulation of SA1, and in both SA1 and SMC1 ChIPs (red and green bars, respectively). We do not believe that there is more cohesin-SA1 at these positions in the siSA1 treated cells. At best, the same amount of cohesin-SA1 is present in control cells and siSA1 treated cells, but it is more efficiently captured by the antibody when the overall levels of SA1 are decreased.

**Fig. S5.**
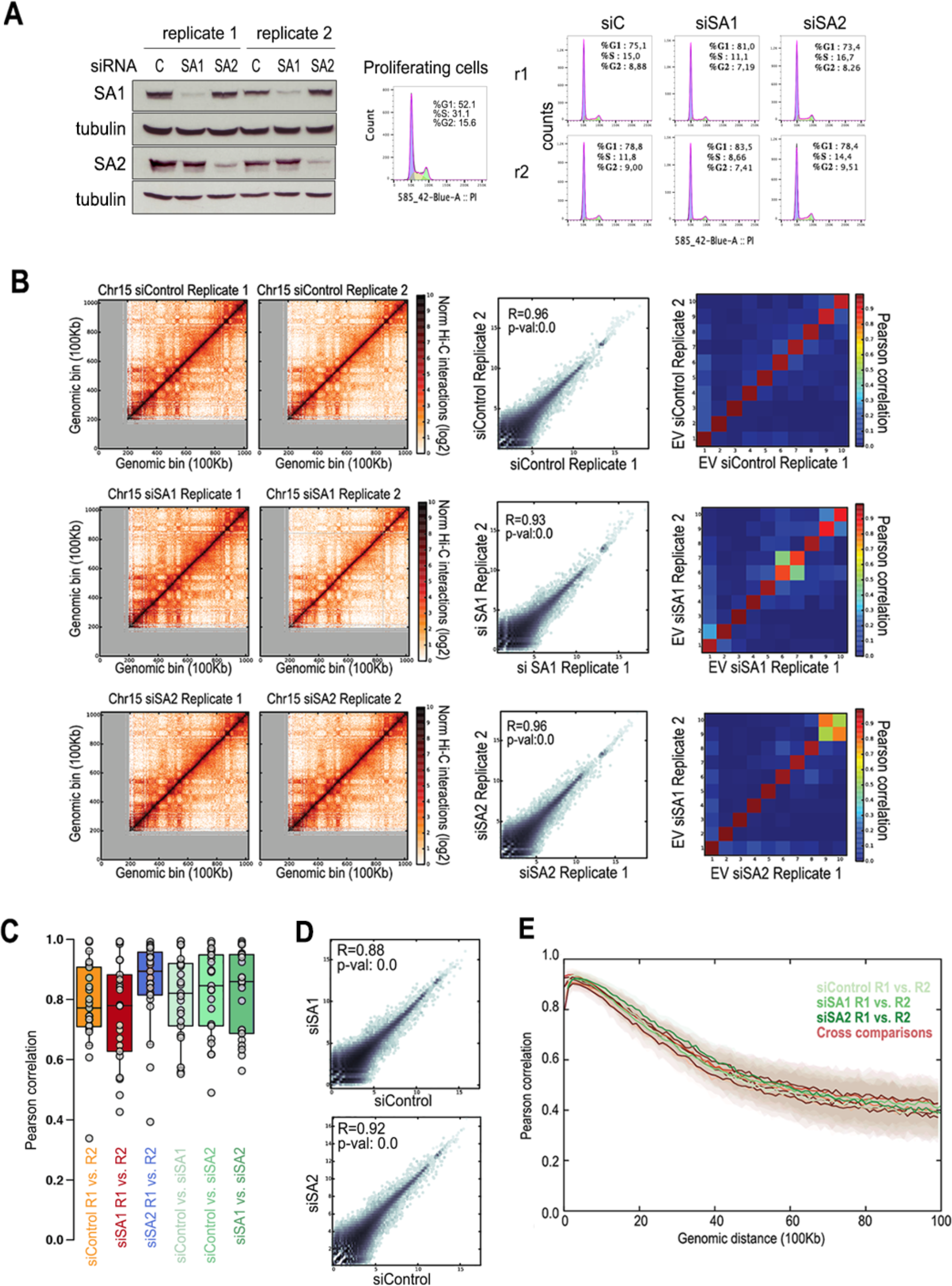
Hi-C analyses in MCF10A cells after depletion of SA1 or SA2. (A) MCF10A cells depleted from SA1 or SA2 by RNAi were arrested in G1 by contact inhibition. Left, immunoblot analysis of the remaining levels of cohesin 72h after transfection. Right, cell cycle profiles of MCF10A proliferating cells and of each of the samples used for Hi-C analysis (two replicas, r1 and r2, per condition). (B) Hi-C normalized interaction maps for chromosome 15 at 100Kb resolution. Left, contact maps for r1 and r2 for each condition (siControl, siSA1 and siSA2). Middle, correlation between r1 and r2 of each condition of all data in the normalized interaction map for chromosome 15. Right, correlation between the Eigen values for the interaction maps for chromosome 15. (C) Box plots of the distribution of Pearson correlation coefficients between Hi-C normalized interactions for all 23 chromosomes. Comparison of Hi-C maps between replicas of each condition and across experiments are shown. (D) Correlation between the siSA1 (top) and siSA2 (bottom) versus siControl for all data in the normalized interaction map for chromosome 15 (E) Correlation between replicates for the Hi-C interaction maps across different genomic distances (from 1 bin to 100 bins of 100Kb).

**Fig. S6.**
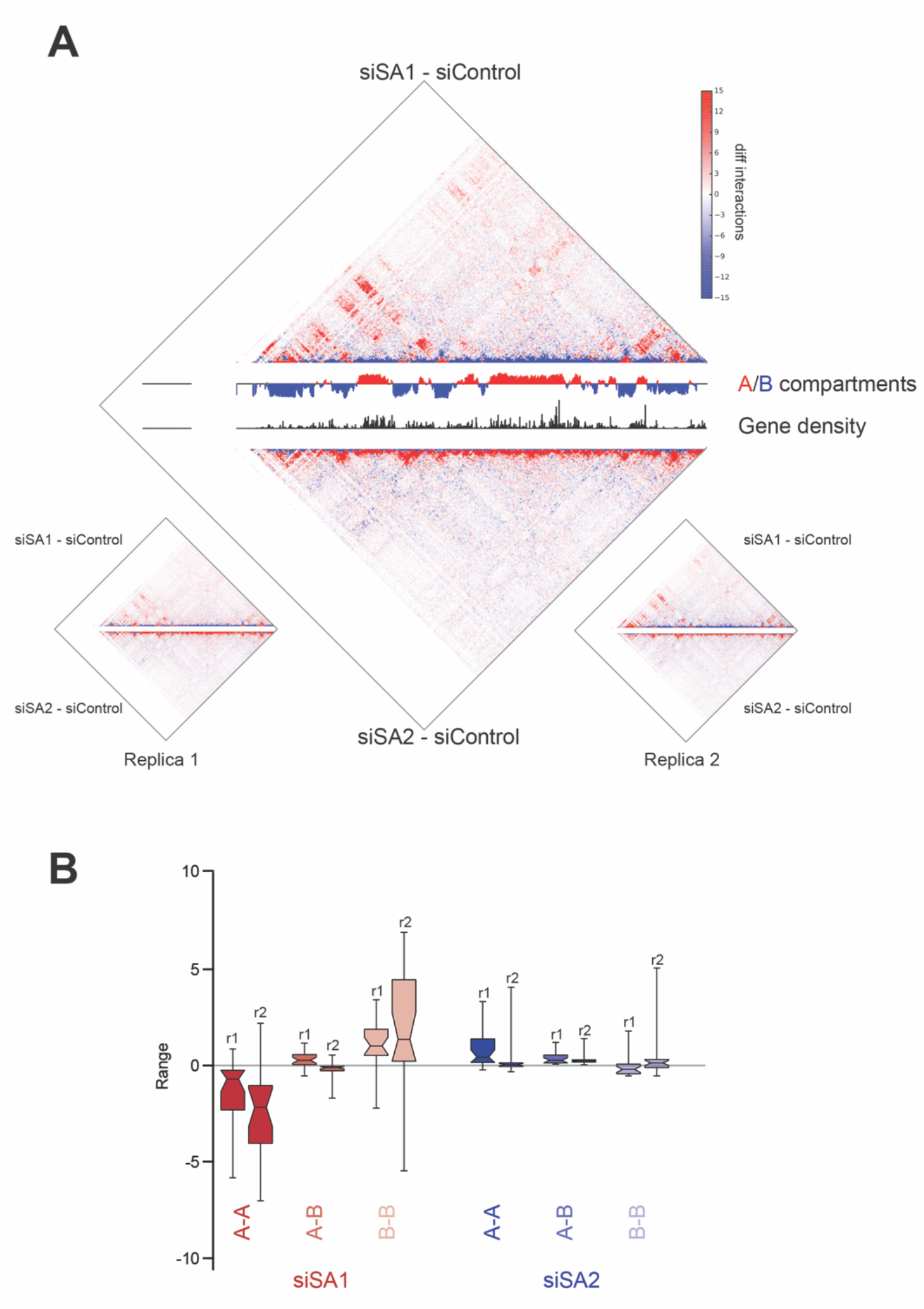
Chromosomal interactions are differently affected by depletion of SA1 or SA2. (A) Chromosome 15 differential interaction map at 100Kb resolution of siSA1 (top)/siSA2 (down) versus siControl for the merged dataset. The panel is complemented by tracks of A/B compartment assignment based on PC1 Eigen values as well as gene density across the genome (middle of the interaction matrices). Replicate experiments are shown on the left for replica 1 and on the right for replica 2 with the same representation. (B) Genome-wide box-plot of chromosome differential interactions separated by replica and A-A, A-B and B-B contacts.

## Tables S1-S7 are provided in a separate Excel file

**Table S1.** ChIP-seq data.

**Table S2.** Proteomic analyses of immunoprecipitates obtained with SA1 and SA2 antibodies in (A) MCF10A cell extracts and (B) HeLa cell nuclear extracts.

**Table S3.** Differentially expressed genes in MCF10A cells after siSA1 treatment

**Table S4.** Differentially expressed genes in MCF10A cells after siSA2 treatment

**Table S5.** Differentially expressed genes in MCF10A cells after siCTCF treatment

**Table S6.** Hi-C data

**Table S7.** Primer sequences

